# The transcriptional basis of quantitative behavioral variation

**DOI:** 10.1101/108266

**Authors:** Kyle M. Benowitz, Elizabeth C. McKinney, Christopher B. Cunningham, Allen J. Moore

## Abstract

What causes individuals to produce quantitatively different phenotypes? While substantial research has focused on the allelic changes that affect phenotype, we know less about how gene expression accompanies variable phenotypes. Here, we investigate the transcriptional basis of variation in parental provisioning using two species of burying beetle, *Nicrophorus orbicollis* and *Nicrophorus vespilloides*. Specifically, we used RNA-seq to compare the transcriptomes of parents that provided high amounts of provisioning behavior versus low amounts in males and females of each species. We found that there were no overarching transcriptional patterns that distinguish high from low caring parents, and no informative transcripts that displayed particularly large expression differences in females or males. However, we did find more subtle gene expression changes between high and low provisioning parents that are consistent across sexes as well as between the two species. Furthermore, we show that transcripts previously implicated in transitioning into parental care in *N. vespilloides* had high variance in the levels of transcription and were unusually likely to display differential expression between high and low provisioning parents. Thus, quantitative behavioral variation appears to reflect many transcriptional differences of small effect. We show that nuanced regulation of the same gene products that are required for the transition of one behavioral state to another are also those influencing variation within a behavioral state.

**Author Summary:** Burying beetles in the genus *Nicrophorus* breed on vertebrate carcasses and provide advanced parental care to their offspring by regurgitating partially digested flesh. However, all adult beetles do not uniformly express this trait. Some provide a large amount of parenting to their offspring, and some only a little. Here, we investigate the genetic causes of why some *Nicrophorus* beetles feed their offspring more than others. We demonstrate that this difference is likely caused by many small changes in gene expression, rather than a few genes that have major effects. We also find that some of the same genes that help to turn on parental care behavior in burying beetles also seem to play a role in determining how much care a beetle gives. These results provide new angles on longstanding questions about the complexity of the mechanisms that underlie quantitative variation in populations.

## Introduction

Understanding the map between genotype and complex phenotype is one of the most important and challenging goals of biology [1–2]. To this end, a great deal of effort has been placed on finding the alleles responsible for producing quantitative variation, through QTL and GWAS methodologies. This avenue of research has been instrumental in uncovering the role of allelic variation in producing heritable quantitative phenotypes [3]. However, intermediate between a variant allele and the quantitative effect it produces must lie some variable mechanistic process [4]. In other words, the mechanism linking genotypic to phenotypic variation must have the potential to be subtly yet reliably altered. Understanding of this kind of mechanistic variation has lagged behind understanding of allelic variation.

Though many biological processes could serve as links from allele to phenotype, one potentially strong candidate is variation in gene expression [5]. If transcription is in fact a mediator of phenotypic effects, then two things must be true: allelic variation must influence transcriptional variation, and transcriptional variation must influence phenotypic variation. Through research mapping QTLs for gene expression phenotypes (eQTL), evidence for the first of these is rapidly mounting [5]. However, empirical work addressing the latter is less common. Research searching for quantitative trait transcripts (QTT; [6]) in *Drosophila* [6–7], human disease [8–9], and crops [10] has suggested that many genes are transcriptionally linked to phenotype [1]. Furthermore, work in *Drosophila* has suggested that predicting candidate transcripts with phenotypic associations may be relatively difficult [11]. However, to date, QTT-like studies have only been performed in a handful of organisms, and none in natural populations, limiting the evolutionary conclusions that can be drawn. Here we begin to address this void by examining how transcriptional variation influences behavioral variation in a complex quantitative behavior, parental provisioning of food by regurgitation to begging offspring.

Behavioral phenotypes can be among the most complex traits found in animals. Behavior is typically highly quantitative in nature, and is produced variably but reliably in response to environmental conditions [12]. As such, behaviors are expected to be underpinned by an especially complicated set of genetic influences [13]. Furthermore, the genetic basis of behavior can be logically split into two processes: first, a process which causes an individual to produce a qualitative change in behavioral state; and second, a process which causes an individual to produce a specific quantitative value of the new behavior. Most studies of transcription and behavior have focused on the first process (reviewed in [14]). For example, in honeybees, numerous investigations have examined the transcriptional basis of the transition between nursing behavior and foraging behavior [15–18], but none have investigated how transcription generates any phenotypic variation within either the nursing or foraging state. A handful of studies investigating the second process (QTT for behavior) have been performed in zebrafish [19–20], and have ascribed a strong and multigenic transcriptional basis to behavioral variation. However, it remains unclear whether the genes that control transitions between behaviors are also involved in producing variation within those same behaviors.

In this study we examine how variation in transcription is associated with variation in parental provisioning behavior in burying beetles (*Nicrophorus* spp.; [21–23]). Recent studies have investigated the transcriptional underpinnings of transitions between behavioral states, such as solitary and parenting, and other parental care behavrios [24–28]. Parenting and specifically the act of provisioning food by regurgitation to begging offspring is an ideal phenotype for contrasting the mechanisms causing behavioral transitions versus within-state variation because it is quantifiable, predictable, and variable [29]. Furthermore, parental care behavior in *Nicrophorus* is known to have a heritable genetic basis [30].

We predicted that quantitative gene expression is related to the quantitative expression of provisioning behavior in *Nicrophorus*. To examine this prediction, we performed RNA-seq comparing gene expression of 20 individuals *a priori* classified as having high or low levels of provisioning behavior based on extensive behavioral observation [29]. Furthermore, we repeated this experiment four times, performing separate RNA-seq experiments in males and females of two species, *N. vespilloides* and *N. orbicollis*. We broadly expected to find many transcripts associated with variation in provisioning behavior in each of the four experiments. Given the strong behavioral [29] but imperfect genetic [27] similarity between sexes, we expected moderate but not complete overlap in the transcripts underlying variation in provisioning between males and females. We predicted we would find even less overlap between species, given the nuanced behavioral differences between them [29, 31]. Additionally, we predicted that the transcripts related to quantitative expression within parenting behavior should be different than those related to transitions between parental and non-parental states in *N. vespilloides* [27]. Our results show that very nuanced changes in gene expression are associated with provisioning behavior and are shared to a degree across sexes and species. Contrary to our predictions, however, we found that the transcripts involved in producing changes between behavioral states are more likely than random to be involved in within-state variation in provisioning behavior.

## Results and Discussion

### Whole genome analysis of parental variation

We first examined whether individual’s overall transcriptomic profile predicted its parental provisioning phenotype. Therefore, we used hierarchical clustering to group all 20 samples in each of the four RNA-seq experiments per the similarity of their genome-wide expression profile. In none of our four experiments did samples of similar phenotype group together (S1–S4 Fig.), and, there is no obvious overall transcriptional signature of being a high or low provisioning parent. We next examined two possibilities: that variation in provisioning behavior is either associated with a few genes of large effect or with many genes of small effect.

To test the first of these possibilities, that variation in provisioning is associated with a few large effect transcripts, we performed standard analysis for differential expression (DE) in edgeR. DE analysis turned up a small number of significantly DE transcripts in three of our four RNA-seq experiments but no obvious patterns (S1 Table). Consistent with Parker et al. [27], more variation in transcription was associated with females but there was no overlap between the species. In males, there was only one transcript that was significantly differentially expressed in male *N. orbicollis* and none in *N. vespilloides* associated with variation in parenting. Because uniparental provisioning behavior is remarkably similar between sexes at both the phenotypic [29] and molecular [27] levels, we expected the DE transcripts found in males and females of a species to at least partially replicate each other. Thus, there is no suggestion that these transcripts are evolutionarily associated with variation in provisioning behavior.

A lack of statistically significant effects does not eliminate the possibility that many genes of small effect contribute to variation in provisioning. Standard analyses of differential expression, as performed above, often do a poor job of finding transcripts with small expression differences between treatments because of multiple testing across the entire transcriptome [32]. Therefore, to test whether such subtle differences might be associated with behavioral phenotype, we asked whether broader patterns of differential expression were similar across the four experiments. To answer this question, we used Gene Set Enrichment Analysis (GSEA; [33]) to assess whether the top DE genes from the standard analysis above displayed unusual patterns of DE in the other three RNA-seq datasets. This should only be the case if two things are true: that there are real transcriptional effects associated with provisioning behavior, and that those effects were at least partially shared across sexes and/or species. Because females have more robust gene expression changes during parenting [27] we used the most DE transcripts from females of each species as a set of “predictor” transcripts, and following Subramanian et al. [33] we *a priori* restricted the analysis to the top 100 transcripts. The top 100 transcripts from *N. vespilloides* females showed a high degree of DE in *N. vespilloides* males (E = 0.582, p = 0.0393; Fig 1A), no unusual DE in *N. orbicollis* females (E = 0.506, p = 0.2944; Fig 1B), and a moderate degree of DE in *N. orbicollis* males (E = 0.578, p = 0.0605; Fig 1C). The top 100 genes from *N. orbicollis* females showed a moderate degree of DE in *N. orbicollis* males (E = 0.625, p = 0.0775; Fig 2A), significant DE in *N. vespilloides* females (E = 0.628, p = 0.0112; Fig 2B), and no unusual DE in *N. vespilloides* males (E = 0.444, p = 0.4294; Fig 2C).

**Fig. 1.**
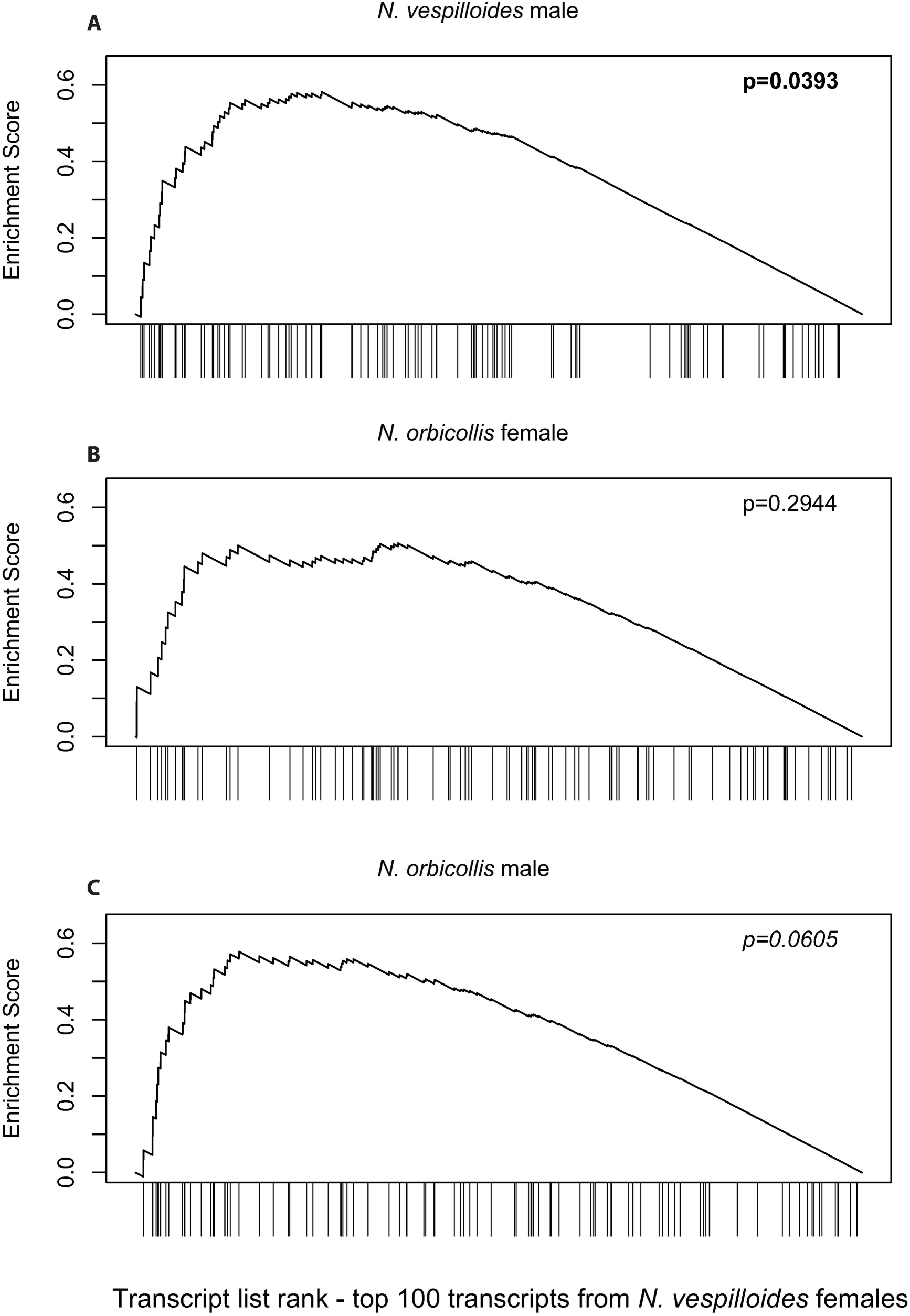
Gene set enrichment analysis of transcripts predicted from *N. vespilloides* females. The top 100 differentially expressed transcripts from *N. vespilloides* females are (**A**) significantly DE in *N. vespilloides* males, (**B**) not DE in *N. orbicollis* females, and (**C**) moderately DE in *N. orbicollis* males.

**Fig 2.**
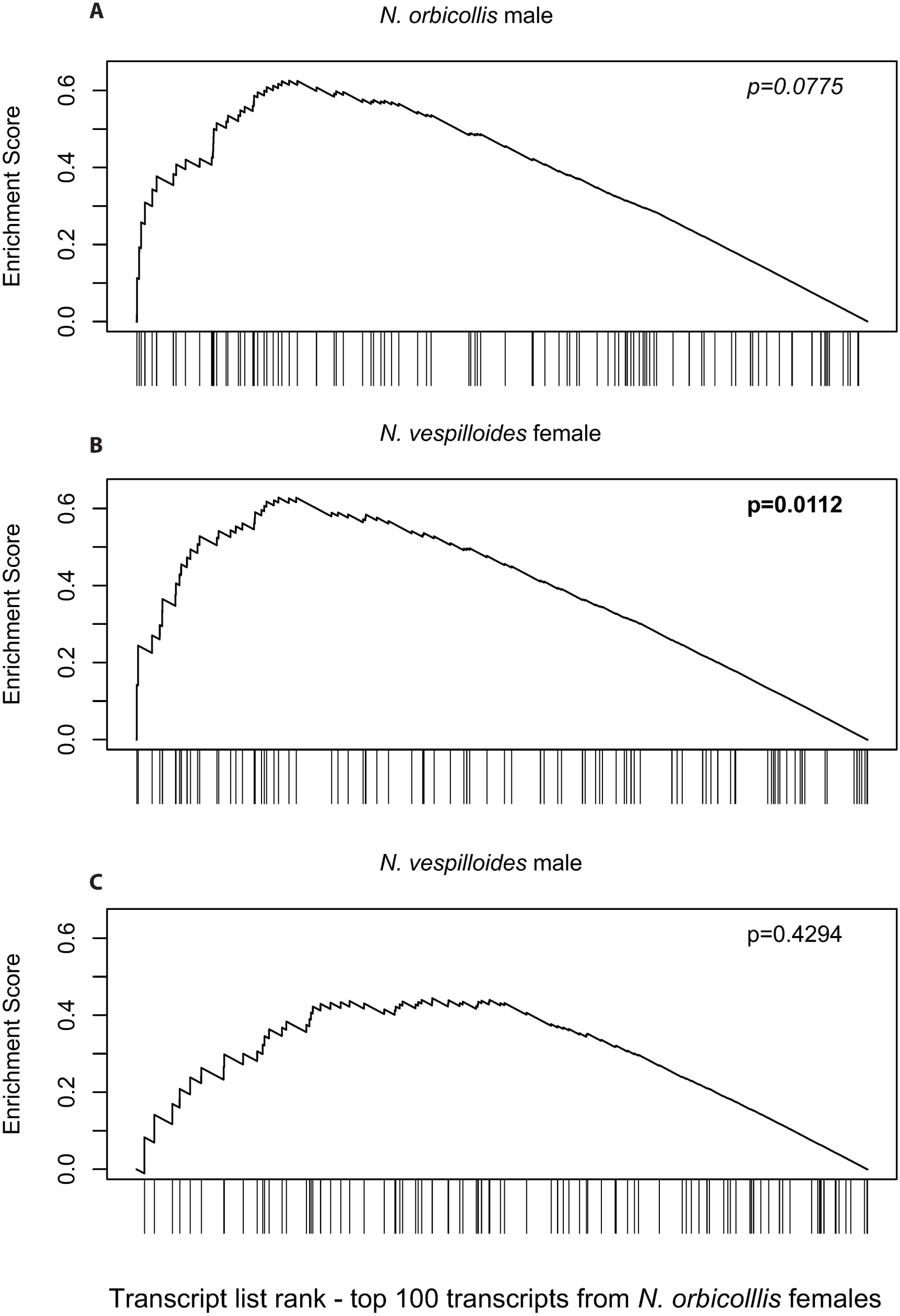
Gene set enrichment analysis of transcripts predicted from *N. orbicollis* females. The top 100 differentially expressed transcripts from *N. orbicollis* females are (**A**) moderately DE in *N. orbicollis* males, (**B**) significantly DE in *N. vespilloides* females, and (**C**) not DE in *N. vespilloides* males.

Our data largely recapitulate the results of other QTT-like studies, which typically demonstrate hundreds of transcripts with significant associations with phenotype [1]. It is unclear why we saw weak statistical associations between phenotype and individual transcripts, though that may simply be a consequence of the messiness of parental care as a phenotype with low but non-zero heritability [30], moderate repeatability [29], or the inability of RNA-seq to measure potentially important classes of transcripts such as neuropeptides [34]. Despite this, our evidence suggests that the transcriptomic basis of variation in *Nicrophorus* provisioning behavior is multifaceted yet extremely subtle, with many genes showing slight associations with phenotype.

This result provides a novel perspective on the longstanding question of whether phenotypic variation is underpinned by a few genes of large effect or many genes of small effect [35]. Though our RNA-seq approach cannot directly answer the question of how many loci are responsible for producing phenotypic variation, it is suggestive of that many variables might be involved. eQTL studies have shown that gene expression phenotypes are likely influenced by many allelic variants [5]. Our results in turn suggest that many gene expression phenotypes are associated with behavioral variation. Therefore, it is likely that there are a large number of allelic variants with the potential to affect provisioning behavior in *Nicrophorus* beetles. Furthermore, it is clear from our data that there are no genes whose transcription has major effects on phenotype. Thus, our results provide continued support for an infinitesimal model of quantitative inheritance [36–37], with many genes contributing an individually undetectable effect on phenotypic expression.

### Analysis of an *a priori* hypothesized gene set

We next asked whether the genes involved in transitioning into parental care behavior are also likely to be involved in generating quantitative variance in provisioning behavior. Parker et al. [27] identified a set of 867 transcripts that were significantly differentially expressed in *N. vespilloides* between parents and non-parents. Using the published transcriptome from Parker et al. [27], we asked whether this gene set displayed unusual patterns of differential expression between high and low provisioning *N. vespilloides*. First, we examined the variability of these genes by comparing the average squared coefficient of variance of genes in this set from that of two random gene sets. We found that, for males (F_2,2250_ = 31.595, p < 0.0001) as well as females (F_2,2250_ = 22.337, p < 0.0001), the caring gene set displays higher variability between samples than random genes (Fig. 3). Next, to see if this high level of variation was related to parental behavior, we used the same GSEA methodology as above. We found that these caring genes were significantly DE in males (E = 0.569, p = 0.0263; Fig 4A) and moderately DE in females (E = 0.570, p = 0.0898; Fig 4B). Thus, it appears that a subset of the genes required to turn on parenting behavior are also modified further to influence the quantitative expression of that same behavior.

**Fig 3.**
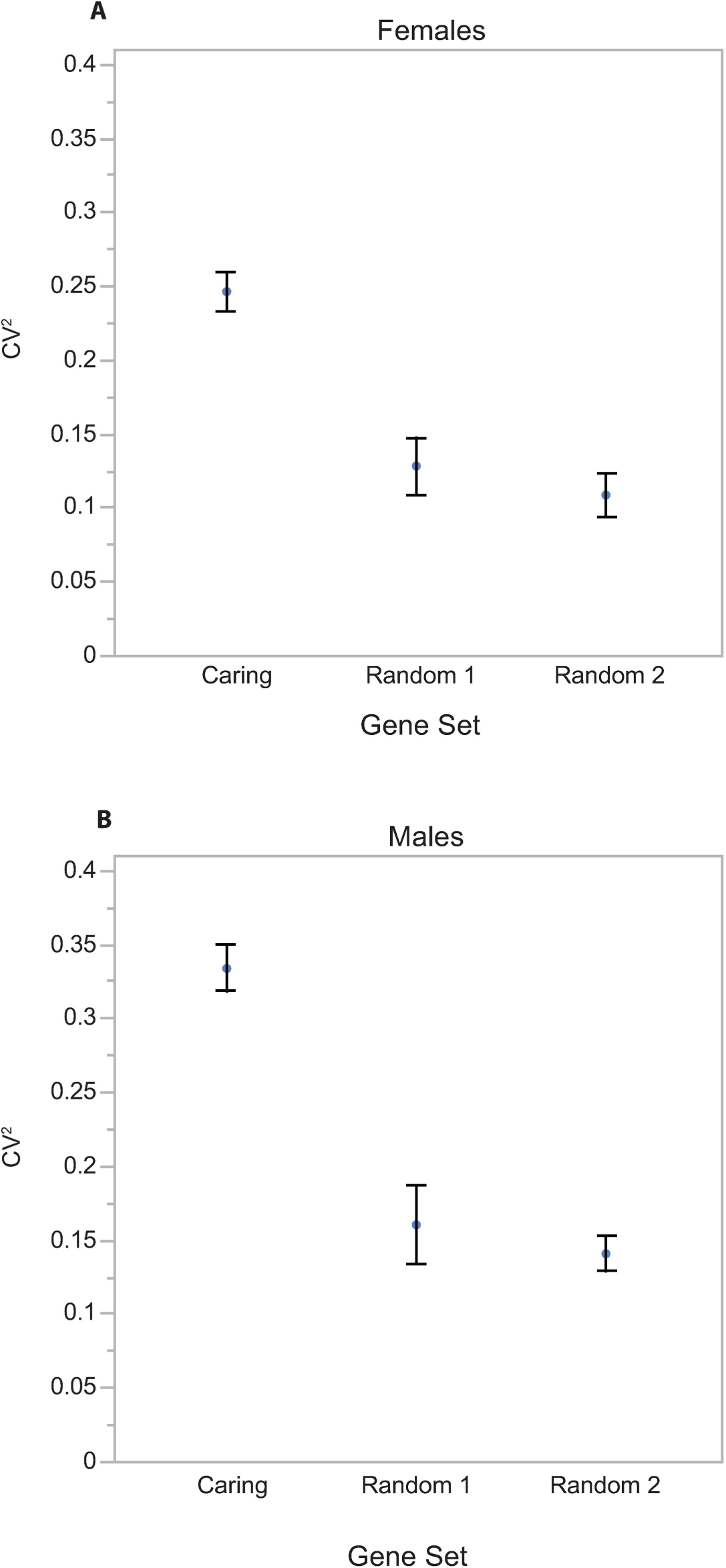
The variance of transcripts implicated in caring versus random genes. Transcripts differentially expressed between caring and non-caring *N. vespilloides* (Parker et al. 2015) show higher variance in our data set than random genes in both (**A**) females and (**B**) males.

**Fig 4.**
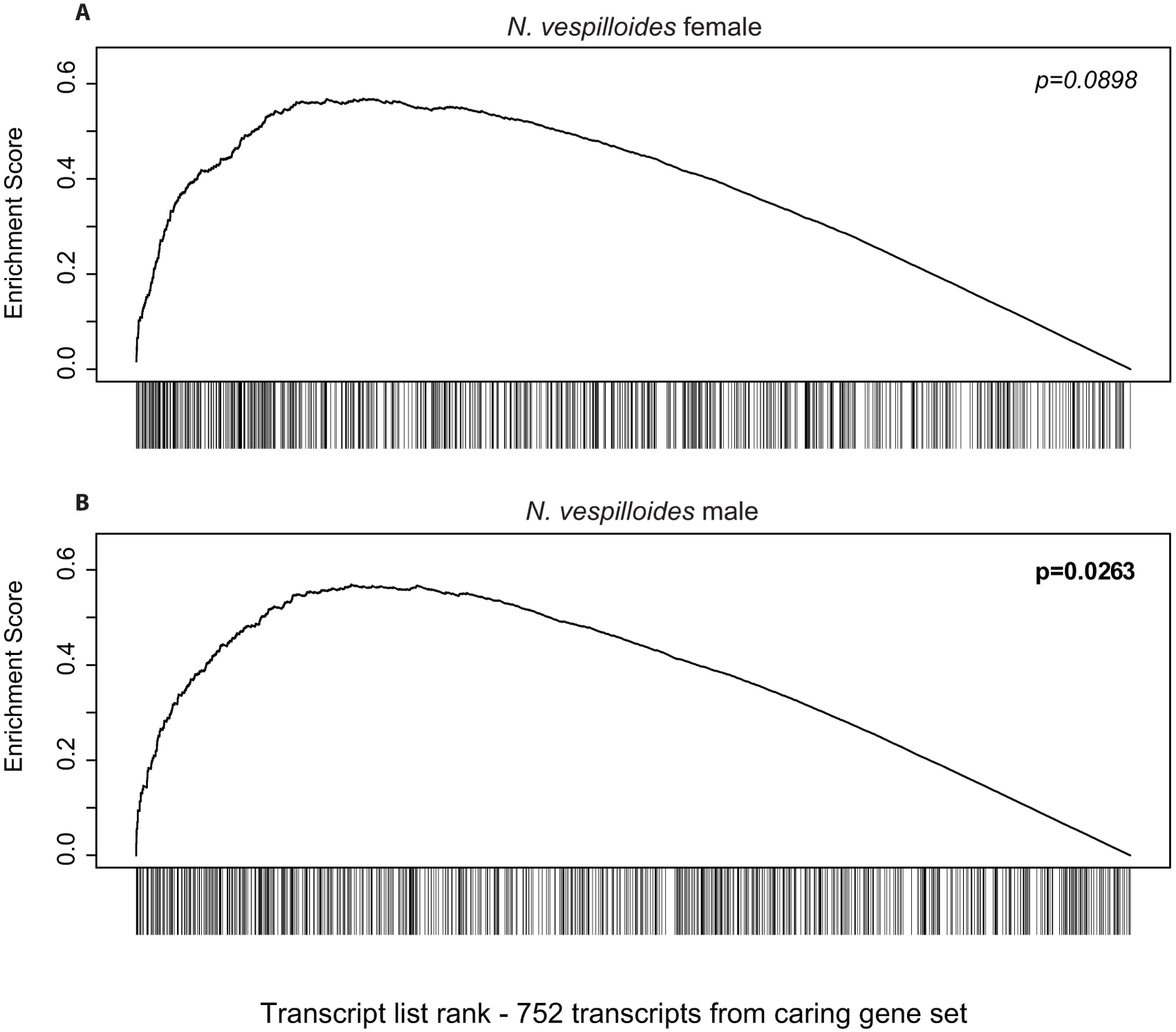
Gene set enrichment analysis of transcripts previously implicated in caring. The 752 transcripts found in our data set that were differentially expressed between caring and non-caring *N. vespilloides* from Parker et al. (2015) are (**A**) moderately DE in *N. vespilloides* females, and (**B**) significantly DE in *N. vespilloides* males.

This result is unexpected when considered in light of morphological developmental biology. The production of behavior is often presented as analogous to development of a morphological structure; there is a mechanism which determines its identity, and then a mechanism to determine its elaboration and final phenotype. For morphological traits, these processes are usually thought to be genetically distinct. In insects, for example, the identities of appendages are specified by Hox transcription factors [38], but the actual process of molding the particular characters of each appendage are largely carried out by other genetic pathways such as WNT [39], Notch [40] and others (but see [41]). The fact that we observe a different pattern here suggests a fundamental difference between the way morphological traits and behavioral traits are regulated. This may be further related to the relatively high flexibility of behavior; if the genes involved in producing behaviors are less constrained than their morphological counterparts, they may be able to produce a broader range of phenotypes in response to environmental variation.

## Conclusions

We present three fundamental insights into the transcriptional architecture of variation in behavior, using parental care in *Nicrophorus* beetles as our model. First, many transcripts appear to affect variation in parenting behavior, each with subtle effects. Second, the mechanisms governing variation in parenting share some commonality across both sexes and species. Third, the transcripts involved in producing transitions into and out of parenting behavior are likely to be involved in producing variation within parenting behavior. Together, these results not only provide basic mechanistic information on the construction of specific behavioral phenotypes, but may also lead to insight as to why behavior is uniquely flexible and evolvable.

## Materials and Methods

### Insect collection and behavioral observation

We collected *Nicrophorus vespilloides* and *Nicrophorus orbicollis* from Cornwall, UK and Georgia, USA, respectively, and maintained outbred laboratory colonies as described in Benowitz et al. [42]. Detailed methods of behavioral experimentation are described in Benowitz et al. [29]. Briefly, we made concurrent observations of female *N. vespilloides* (n = 57) and *N. orbicollis* (n = 61) during the hours of peak parental care, approximately one day after larval hatching. We scan-sampled each parent 80 times over an eight-hour period, recording each time she was observed having mouth-to-mouth contact with larvae. We then repeated the experiment under the same conditions for males of *N. vespilloides* (n = 79) and *N. orbicollis* (n =78). After we finished observations of each parent, we removed their heads, flash-froze them in liquid nitrogen, and stored them at −80°C.

### Sample collection, preparation, and RNA sequencing

After we completed all behavioral observations, we selected the 10 highest and 10 lowest caring parents from each species and sex for our RNA-seq experiment. We extracted RNA from these 80 individuals by first homogenizing them in liquid nitrogen and Qiazol (Qiagen, Venlo, Netherlands), then adding chloroform [26]. After this step, extractions followed standard protocols from a Qiagen RNeasy Lipid kit. RNA libraries were then prepared using a TruSeq Stranded RNA LT Kit (Illumina, San Diego, CA) with one-third of the normal reaction volume [43]. Samples were then sequenced on an Illumina NextSeq Mid Output Flow Cell at the Georgia Genomics Facility (GGF), generating paired-end 75 base pair reads.

### *N. orbicollis* genome sequencing and assembly

In order to produce comparable transcriptomic analyses between species, we decided to sequence the *N. orbicollis* genome to match the recently published *N. vespilloides* genome [44]. For genome sequencing, we chose a single larva that was the product of two generations of inbreeding. We extracted its DNA using a sodium dodecyl sulfate (SDS) lysis buffer [45] and a phenol chloroform extraction. Two libraries, one 350 bp PCR-free library and one 6.5kb mate-pair library were made using Illumina manufacturer’s instructions, and sequenced on an Illumina NextSeq Mid Output Flow Cell at GGF generating paired-end 75 base pair reads. We next trimmed reads for quality using Trimmomatic 0.32 [46], removing 4bp windows with average quality less than 15 and removing reads under 25bp. We assembled the trimmed reads with Platanus [47] under the same parameters as for *N. vespilloides* [44]. We then used DeconSeq [48] to remove any possible contaminants from the genome assembly. Genome assembly was 193 mb, with an N50 of 178.7 kb. BUSCO analysis [49] showed the assembly had complete sequence for 82.9% of conserved insect orthologs, and partial sequence for 88.6%.

### Transcriptome assembly and gene expression analysis

We trimmed all RNA reads for quality using Trimmomatic 0.32 as above [46] and mapped them to their respective reference genome using Tophat 2 [50]. We then assembled reads for each species into transcripts using Cufflinks [51], and taking the longest isoform provided. We then remapped reads to the assembled and reduced transcriptome and generated gene level counts using RSEM [52]. Descriptive statistics for each assembled transcriptome are provided in Table S2.

Next, we used edgeR [53] to filter transcripts by abundance and perform normalization. Normalized counts were used to perform hierarchical clustering analysis using the R package hclust under default parameters. We then performed differential expression (DE) analysis in edgeR between high and low caring parents. We analyzed each species and sex separately, thus creating four separate DE analyses, assessing significance after multiple testing using FDR [54].

### Gene set enrichment analysis

We used Gene Set Enrichment Analysis (GSEA) to examine similarity between our four independent datasets. Following Subramanian et al. [33], we selected the top 100 differentially expressed genes from one dataset (*N. vespilloides* females) to serve as a predictive set of genes. To examine this set of genes in another dataset, we calculated a running-sum statistic going down the entire list of genes (ordered from most DE to least DE). This statistic increases each time a gene in the set is encountered (weighted by the amount of DE) and decreases by a constant amount each time a gene outside of the set is encountered. The test statistic, or E-score, is the global maximum of this running sum statistic. The significance of the E-score was calculated by comparing it against 10,000 E-scores generated from gene lists with randomized phenotypic permutations. Therefore, this analysis asks whether the predictive gene set displays more DE using the real phenotypic information than random phenotypic information, a conservative GSEA methodology [33,55]. For the set of 100 genes taken from *N. vespilloides* females, we performed this analysis in *N. vespilloides* males, *N. orbicollis* females, and *N. orbicollis* males. We also repeated the same analysis using the top 100 differentially expressed genes from *N. orbicollis* females. This predictive gene set was analyzed in *N. orbicollis* males, *N. vespilloides* females, and *N. vespilloides* males. We performed all GSEA analysis in R using homemade scripts.

### Mapping and differential expression analysis using previously published transcriptome

In order to compare our results directly to those of Parker et al. [27] we mapped our *N. vespilloides* RNA-seq reads directly to their published transcriptome using Tophat2 [50] and generated read counts using RSEM [52]. Gene level counts were then filtered and normalized in edgeR as above. We then specifically examined the 867 (752 after abundance filtering) genes differentially expressed during parenting in *N. vespilloides* [27]. First, we examined whether these genes displayed greater variability during parenting than non-caring genes by comparing the average squared coefficient of variance (CV^2^) to that of two random gene sets of equal size and average abundance using analysis of variance. We then performed GSEA using the same methodology as above to examine whether these genes displayed unusual patterns of differential expression between parents with high versus low provisioning phenotypes.

## Acknowledgments

B. Hsu and V. Anderson assisted with *Nicrophorus* colony maintenance. B. Schmitz, D. Neumann, N. Rohr, K. Sandlin, J. Wagner, and M. Alabady assisted with library preparation and Illumina sequencing. D. Hall, K. Ross, T. Linksvayer, A. Wiberg, D. Parker, and M. Ritchie provided helpful discussions and suggestions for statistical analysis. T. Moore provided valuable comments on the manuscript. Sequence data associated with this project is available through NCBI (Bioproject # PRJNA371654). Behavioral data is available through Dryad (doi:10.5061/dryad.25rm2).

## Figure Captions

**S1 Fig. Hierarchical clustering of *N. orbicollis* female samples by transcriptional profile.**

**S2 Fig. Hierarchical clustering of *N. orbicollis* male samples by transcriptional profile.**

**S3 Fig. Hierarchical clustering of **N. vespilloides** female samples by transcriptional profile.**

**S4 Fig. Hierarchical clustering of **N. vespilloides** male samples by transcriptional profile.**

**S1 Table. Transcripts significantly differentially expressed between high and low provisioning parents.** A list of the significantly DE transcript IDs from each experiment and their blast homologies. Asterisks next transcripts from **N. orbicollis** females indicate transcripts whose differential expression is explained by extremely high expression values in the same 3 (out of 20) individuals, and therefore may be outliers.

